# Cooperative Learning with Penalized Linear Mixed-Effects Models for High-Dimensional Clustered Multiview Data

**DOI:** 10.64898/2026.09.01.748742

**Authors:** Shunsuke Yoshimura, Mariko Takagishi, Kensuke Tanioka

## Abstract

In biomedical research, multiple types of high-dimensional data, such as genomic, transcriptomic, proteomic, and metabolomic data, are increasingly collected from the same subjects. Integrating these multiple data views can improve prediction by exploiting shared or complementary information across the views. Cooperative learning provides an agreement-based framework for multiview supervised learning by encouraging predictions obtained from individual views to be similar. However, the original framework assumes independent observations and therefore does not account for clustered structures, such as repeated measurements obtained from the same subject. To address this limitation, we propose Cooperative Learning with a penalized Linear Mixed Model (CL-pLMM) for high-dimensional multiview data with a clustered structure. CL-pLMM replaces the ordinary prediction loss in cooperative learning with a covariance-weighted loss that accounts for within-cluster dependence, while retaining the agreement penalty between views and a Lasso penalty for variable selection. We further show that its objective function can be represented as a penalized linear mixed-effects model applied to augmented data, allowing existing estimation procedures to be used. The performance of CL-pLMM is evaluated through simulation studies under various signal and dependence settings and an application to longitudinal proteomic and metabolomic data for predicting the time to spontaneous labor.

## 1 Introduction

In recent years, it has become increasingly common to collect multiple types of omics data, such as genomics, transcriptomics, proteomics, and metabolomics, from the same subjects. Integrating multiple data views may improve outcome prediction by leveraging information that cannot be captured by a single view alone. Basic approaches to multiview data integration include early fusion and late fusion. Early fusion combines multiple views before constructing a prediction model, whereas late fusion constructs prediction models separately for each view and subsequently combines their predictions. However, these simple integration approaches may not fully exploit the relationships among different views.

Ding et al. [1] proposed cooperative learning for multiview supervised learning. Cooperative learning predicts an outcome using multiple views while introducing an agreement penalty that encourages the predictions obtained from the individual views to agree with each other, thereby exploiting predictive information shared across views. Ding et al. [1] applied this method to multi-omics data, including proteomics and metabolomics data. Cooperative learning has subsequently been extended to various settings, including semi-supervised learning [2], interaction modeling [3], heterogeneous multiview data [4], and multi-cohort data integration [5].

In many biomedical and other scientific studies, observations are not necessarily independent but may have a clustered or grouped structure, where multiple observations belong to the same cluster. Such a cluster may, for example, correspond to a subject or experimental unit from which multiple observations are obtained. Longitudinal data, in which the same subjects are repeatedly measured over time, are an important example of such data. Observations within the same cluster generally cannot be regarded as independent, and their correlation therefore needs to be taken into account. Linear mixed-effects models (LMMs) provide a widely used framework for accounting for correlation among observations in grouped or clustered data [6].

For high-dimensional data such as omics data, it is necessary to account for correlation among observations while selecting relevant variables from a large number of predictors. Penalized mixed-effects models have been developed for this purpose. Bhatnagar et al. [7] proposed a penalized linear mixed model and an estimation procedure that introduces a sparsity-inducing penalty on the fixed effects. Their method enables variable selection among high-dimensional predictors while accounting for correlation or relatedness among observations. Various penalized mixed-effects models have subsequently been proposed for high-dimensional longitudinal and clustered data [8], [9], [10]. However, these methods are not designed to integrate multiview information by exploiting agreement between predictions obtained from multiple views, as in the cooperative learning framework of Ding et al. [1].

Methods that combine multi-omics or multiview integration with dependence structures or random effects have also been developed. For example, Bayesian frameworks and related approaches have also been proposed for integrating multi-omics data, including methods for longitudinal or disease-progression data [11], high-dimensional prediction [12], and prediction and classification [13]. A more closely related approach is Bayesian Integrative Analysis and Prediction with mixed effects (BIPmixed) proposed by Neher et al. [14]. Its data integration strategy is based on supervised intermediate fusion, which exploits latent structures shared across multiple views. This strategy differs from the cooperative learning framework of Ding et al. [1], which directly exploits agreement between predictions obtained from the individual views. The agreement-based formulation of cooperative learning provides a relatively simple and straightforward-to-implement strategy for multiview integration.

Accordingly, in this study, we propose Cooperative Learning with a penalized Linear Mixed Model (CL-pLMM), which extends the cooperative learning framework of Ding et al. [1] to clustered multiview data. Specifically, CL-pLMM retains the key feature of cooperative learning by encouraging agreement between predictions obtained from the individual views, while accounting for within-cluster correlation through a mixed-effects covariance structure. It also performs variable selection among high-dimensional predictors using a sparsity-inducing penalty. We evaluate the performance of the CL-pLMM through a simulation study and further apply it to longitudinal multi-omics data analyzed by Ding et al. [1].

## 2 Background Methods

### 2.1 cooperative learning

We first introduce the objective function of cooperative learning, which forms the basis of CL-pLMM, together with its augmented-data representation. Throughout this paper, we focus on multiview data consisting of two data views. Let ***y*** ∈ ℝ^*n*^ denote a centered response vector, and 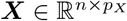 and 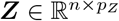 denote centered design matrices for the two data views. Cooperative learning [1] minimizes the following objective function:

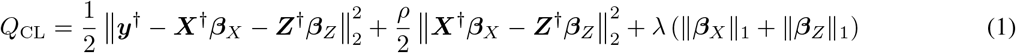

where 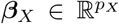 and 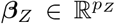 the prediction error based on t, and *λ, ρ >* 0 are tuning parameters. Here, ∥***a***∥_2_ = (***a***^*T*^***a***)^1*/*2^, and 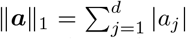 for any vector ***a*** = (*a*_1_, …, *a*_*d*_)^*T*^ ∈ ℝ^*d*^, where ***a***^*T*^ is the transpose of ***a***. The first term in (1) measures the prediction error based on the two views, the second term is an agreement term that encourages the linear predictors obtained from the two views to be similar, and the last term is a Lasso penalty for variable selection.

Define the augmented response vector, augmented design matrix, and coefficient vector as

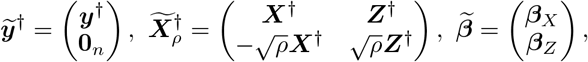

where **0**_*n*_ denotes the zero vector of length *n*. Then, Eq. (1) can be written as

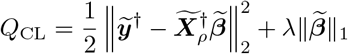

which is equivalent to a Lasso regression applied to the augmented data 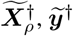 .

### 2.2 Penalized Linear Mixed-Effects Model for Clustered Data

In many applications, observations have a clustered or grouped structure, such that multiple observations belong to the same cluster and may therefore be correlated. Hereafter, we use the term clustered data. A cluster may represent, for example, a subject or an experimental unit from which multiple observations are obtained. LMMs account for dependence among observations within the same cluster through random effects.

Suppose that the data consist of *m* clusters. Let *n*_*k*_ denote the number of observations in cluster *k*, (*k* = 1, …, *m*), and let 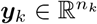denote the centered response vector for the cluster *k*, and let 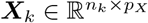 denote the centered design matrix for cluster *k*. Let 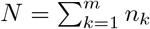 be the total number of observations, and define the stacked response vector and design matrix by 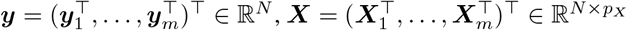 . We consider the linear mixed-effects model with a single random effect [15]

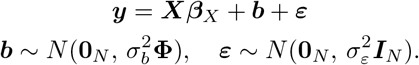

Here, 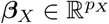 denotes the fixed-effect coefficient vector, ***b*** is the random-effect vector, ***ε*** is the vector of error terms, both of which are random vectors of length *N* . 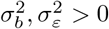 are unknown variance parameters for ***b*** and ***ε***, respectively. The matrix **Φ** ∈ ℝ^*N ×N*^ is a prespecified positive semidefinite matrix representing the dependence structure among the clustered observations. Its specific form may be chosen according to the cluster structure and the application. Let 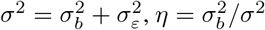 . Then,

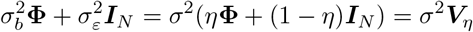

where ***V***_*η*_ = *η***Φ** + (1 − *η*)***I***_*N*_, and ***I***_*N*_ is the *N* × *N* identity matrix. Therefore, we get

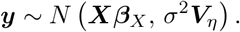

Ignoring terms that do not depend on the model parameters, the negative log-likelihood 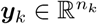 is

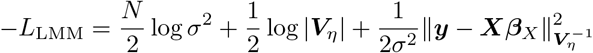

Here, |***B***| denotes the determinant of a square matrix ***B***, and ∥***a***∥_***B***_ = (***a***^*T*^***Ba***)^1*/*2^. The first two terms are likelihood terms associated with estimation of the variance parameters, while the third term is a Mahalanobis-type prediction error that accounts for dependence among observations.

To enable simultaneous variable selection and effect estimation in high-dimensional settings, Bhatnagar et al. [7] considered the corresponding penalized objective function given by

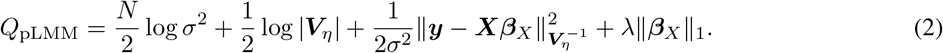

## 3 Proposed CL-pLMM

We consider clustered multiview data comprising two data views, in which multiple observations belong to the same cluster. In addition to exploiting information shared across views, as in standard cooperative learning, dependence among observations within the same cluster needs to be taken into account. We therefore extend cooperative learning so that agreement between the linear predictors from the two views is encouraged while within-cluster correlation is accounted for through a mixed-effects model.

For *k* = 1, …, *m*, let 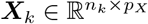 denote the centered response vector for cluster *k*, 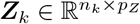 and 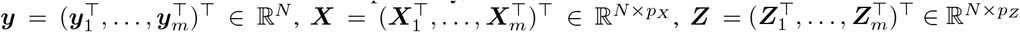 denote the centered design matrices for the first and second data views, respectively, for cluster *k*. Define the stacked response and design matrices as 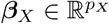 .

To extend cooperative learning to clustered data, we replace the ordinary squared prediction loss in cooperative learning with a covariance-weighted loss that accounts for within-cluster dependence, while retaining the agreement term between the linear predictors from the two views. Based on the cooperative learning objective in Eq. (1) and the penalized mixed-effects objective in Eq. (2), we define the proposed objective function as

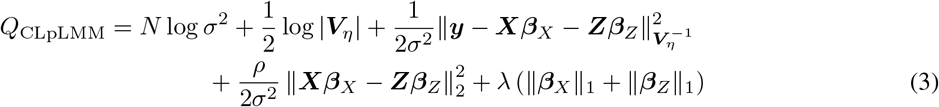

where 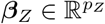 and 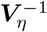 are unknown fixed-effect coefficient vectors for the two data views, and *σ*^2^ *>* 0 and *η* ∈ [0, 1] are unknown variance parameters. *ρ >* 0 and *λ >* 0 are tuning parameters.

The matrix ***V***_*η*_ is defined, as in Section 2.2, by ***V***_*η*_ = *η***Φ** + (1 − *η*)***I***_*N*_ where **Φ** ∈ ℝ^*N ×N*^ is a prespecified positive semidefinite dependence matrix. Specific forms of **Φ** used in the simulation study and the real-data application are described in Sections 4 and 5, respectively.

In Eq. (3), the first two terms are likelihood terms associated with the estimation of the variance parameters. The third term measures the prediction error using the covariance-weighted norm induced by 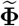 and therefore accounts for within-cluster dependence. The fourth term measures the discrepancy between the linear predictors obtained from the two views using the Euclidean norm, as in standard cooperative learning, and thereby encourages them to be similar. The final term is a Lasso penalty on ***β***_*X*_ and ***β***_*Z*_, which performs variable selection within each view.

To express Eq.( 3) as a penalized linear mixed-model objective, define the augmented response vector, design matrix, coefficient vector, and dependence matrix as

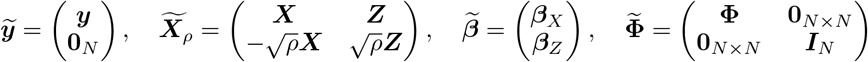

where **0**_*N* × *N*_ denotes the *N* × *N* zero matrix. The upper-left block of 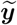 contains **Φ**, which represents the dependence structure among the original clustered observations, whereas the lower-right block is ***I***_*N*_ . Thus, the prediction error is measured using the covariance structure ***V***_*η*_, whereas the discrepancy between the linear predictors from the two views is measured using the Euclidean norm.

For the augmented data, consider

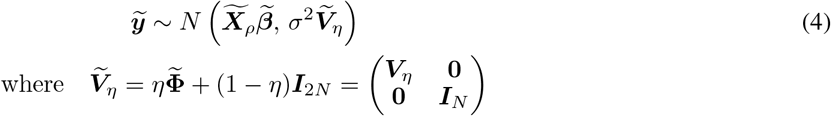

Ignoring terms that do not depend on the model parameters, the negative log-likelihood for 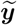 is

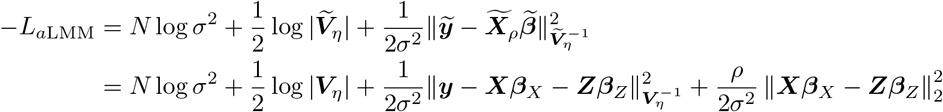

Adding a Lasso penalty on 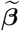 gives

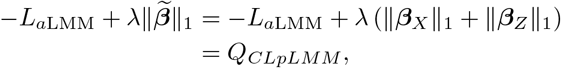

meaning that this yields the function which is identical to the proposed objective function in Eq. (3). Hence, CL-pLMM can be viewed as a penalized linear mixed-effects model applied to the augmented data in Eq. (4). This representation allows the model parameters to be estimated using the same estimation framework as the penalized LMM in Eq. (2). For details of the estimation algorithm for the penalized LMM, see [7].

## 4 Numerical Simulation

### 4.1 Data Generation

We conducted simulation studies to evaluate the performance of the proposed CL-pLMM. Artificial datasets were generated based on the latent factor model of [1], extended to accommodate clustered data. Each dataset consisted of two high-dimensional data views, 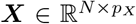 and 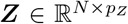, measured for the same *N* observations. The observations were grouped into *m* clusters, with cluster *k* containing *n*_*k*_ observations, so that 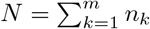.

The data were generated in three steps. First, baseline predictor variables were generated independently for the two views. Second, shared latent factors were added to a subset of variables in both views, thereby inducing associations between the two views and the response. Finally, correlation among observations within the same cluster was introduced through the error structure of the response.

Specifically, for observation *j* = 1, …, *n*_*k*_ in cluster *k* = 1, …, *m*, the baseline predictor variables were generated as

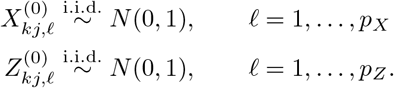

To create informative variables shared across the two views, *p*_*u*_ latent-factor scores were independently generated for each observation as

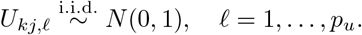

These latent factors were added to the first *p*_*u*_ variables of each view:

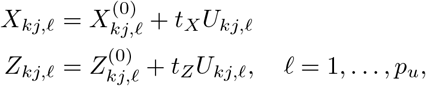

where *t*_*X*_ and *t*_*Z*_ control the strength of the latent-factor contribution to the *X* and *Z* views, respectively. The remaining variables were defined as 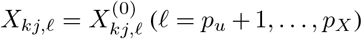 and 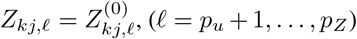 . Thus, only the first *p*_*u*_ variables in each view were associated with the latent factors, whereas the remaining variables were unrelated to the response.

The true signal for observation *j* in cluster *k* was generated from the same latent factors as

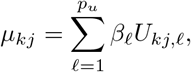

where *β*_*l*_ denotes the contribution of the *l*th latent factor to the response. Consequently, the first *p*_*u*_ variables in each view were indirectly associated with the response through the shared latent factors.

To introduce correlation among observations belonging to the same cluster, the response vector was generated as

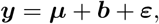

where ***b*** ∼ *N* (**0**_*N*_, *ησ*^2^**Φ**) and ***ε*** ∼ *N* (**0**_*N*_, (1 − *η*)*σ*^2^***I***_*N*_ ). Here, ***b*** induces correlation among observations within the same cluster, whereas ***ε*** represents observation-specific error. For the simulation study, the dependence matrix **Φ** ∈ ℝ^*N ×N*^ was specified as the block-diagonal matrix

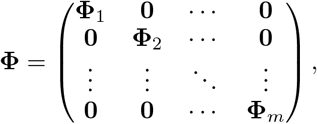

where the block corresponding to cluster *k* was

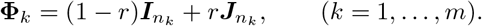

Here, 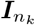 and 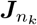 denote the *n*_*k*_ × *n*_*k*_ identity matrix and all-ones matrix, respectively. Thus, the diagonal elements of **Φ**_*k*_ were one, and its off-diagonal elements were *r*. This specification represents an exchangeable dependence structure within each cluster, with observations from different clusters assumed to be uncorrelated.

Unlike Ding et al. [1], who fixed the noise standard deviation to obtain an approximately specified signal-to-noise ratio (SNR), we determined *σ*^2^ separately for each simulated dataset. The SNR was defined as SNR = Var(***µ***)*/σ*^2^, where Var(***µ***) denotes the variance of the true signal across all generated observations.

### 4.2 Simulation Study Design

We set *m* = 2000 clusters and *n*_*k*_ = 5 observations for all *k* = 1, …, *m*, resulting in 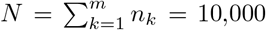 observations in total. The numbers of variables were *p*_*X*_ = *p*_*Z*_ = 500, and *p*_*u*_ = 30 latent factors were used. The latent-factor coefficients were fixed at *β*_*l*_ = 8 for *l* = 1, …, *p*_*u*_.

For each simulation replicate, 40 clusters (200 observations) were randomly assigned to the training set, and the remaining 1960 clusters (9800 observations) were used as the test set. The training–test split was performed at the cluster level so that repeated measurements from the same cluster never appeared in both sets.

We considered three signal configurations based on the simulation settings of [1]: (*t*_*X*_, *t*_*Z*_, SNR) = (2, 2, 1.8), (6, 1, 0.6), and (2, 0, 3.5). These configurations represent, respectively, a setting in which both views contain comparable amounts of signal, a low-SNR setting with an asymmetric signal contribution from the two views, and a setting in which only the ***X*** view contains signal associated with the response.

The variance-component parameter was fixed at *η* = 0.5. For each signal configuration, three levels of within-cluster dependence, *r* ∈ {0, 0.3, 0.6}, were considered. Thus, a total of nine simulation scenarios were examined. Since *η* = 0.5, these values of *r* correspond to within-cluster residual correlations of 0, 0.15, and 0.30, respectively.

Before model fitting, each variable in ***X*** and ***Z*** was standardized using the mean and standard deviation estimated from the training data. The same transformation was then applied to the test data. The response was centered using the mean response in the training data.

We compared ten prediction methods: Separate *X*, Separate *Z*, Early Fusion, Late Fusion, Cooperative Learning, Separate *X*-pLMM, Separate *Z*-pLMM, Early Fusion-pLMM, Late Fusion-pLMM, and CL-pLMM. Separate *X* and Separate *Z* fit Lasso models using only the corresponding view. Early Fusion concatenates ***X*** and ***Z*** and fits a single Lasso model, whereas Late Fusion fits separate Lasso models to the two views and combines their predictions using estimated weights. Cooperative Learning jointly fits the two views using an agreement penalty.

The corresponding pLMM-based methods replace the Lasso model with a penalized linear mixed model that accounts for within-cluster dependence through **Φ**. The pLMM-based methods were estimated using the `ggmix` package in R [7], with CL-pLMM fitted by applying `ggmix` to the augmented data representation shown in Eq. (4). Because `ggmix` includes an intercept by default, the response and predictors were centered before model fitting to limit the effect of the estimated intercept on the agreement term in the augmented formulation.

Regularization parameters were selected using five-fold cross-validation. The folds were constructed at the cluster level so that all repeated measurements from the same cluster were assigned to the same fold. For Cooperative Learning and CL-pLMM, the agreement parameter *ρ* was selected from {0, 0.2, 0.4, 0.6, 0.8, 1}. For Late Fusion and Late Fusion-pLMM, 30% of the training clusters were additionally reserved as a validation set for estimating the weights used to combine predictions from the two views.

Each of the nine simulation scenarios was independently repeated 100 times, with a new dataset generated for every replicate.

### 4.3 Evaluation

Prediction performance was evaluated on the held-out test set with respect to the underlying true signal ***µ***. Let 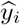 denote the response predicted by each fitted model for the *i*th test observation (*i* = 1, …, *N*_test_), where *N*_test_ is the number of test samples. The test mean squared error (MSE) was calculated as 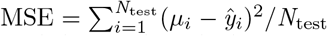 . In addition to prediction performance, the total number of selected variables was recorded for each method.

### 4.4 Results

Fig. 1 shows the test MSE across the nine simulation scenarios. When both views contained comparable amounts of signal, Cooperative Learning achieved lower MSE than the separate, early-fusion, and late-fusion approaches. CL-pLMM achieved the lowest MSE for all three levels of within-cluster dependence. The difference in MSE between Cooperative Learning and CL-pLMM was small when *r* = 0 but increased as *r* increased.

**Figure 1.**
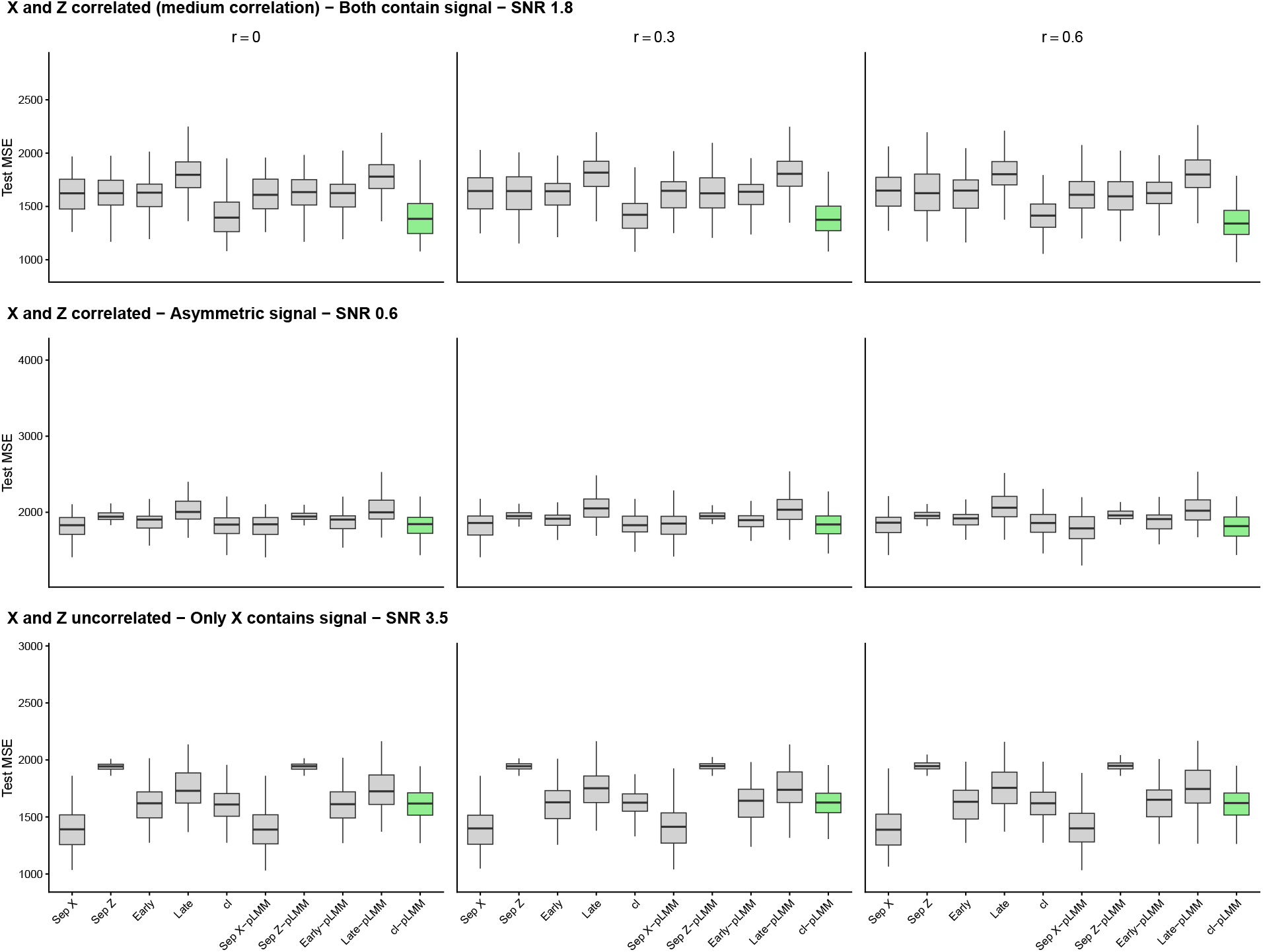
Test MSE for the nine simulation scenarios. Rows correspond to the three cross-view signal configurations, and columns correspond to the within-cluster dependence parameter *r* = 0, 0.3, and 0.6. The horizontal axis indicates the methods compared: “Sep X” and “Sep Z” denote Separate X and Separate Z, respectively; “Early” and “Late” denote Early Fusion and Late Fusion, respectively; and “CL” denotes Cooperative Learning. The suffix “-pLMM” indicates the corresponding method based on a penalized linear mixed model. CL-pLMM is shown in green.

When the signal contribution was asymmetric and the SNR was low, Separate *X* achieved the lowest MSE for *r* = 0 and 0.3, whereas Separate *X*-pLMM achieved the lowest MSE for *r* = 0.6. Although CL-pLMM did not outperform Separate *X* in this setting, it achieved lower MSE than Cooperative Learning when within-cluster dependence was strongest.

When only *X* contained signal, Separate *X* consistently achieved the lowest MSE. The relative performance of the methods was largely unchanged across the three values of *r*.

## 5 Application to Labor Onset Data

### 5.1 Data and analysis objective

In this section, we analyzed the labor onset dataset consisting of pregnant women who eventually experienced spontaneous labor [16]. The objective was to predict the number of days until spontaneous labor, referred to as days to spontaneous labor (DOS), using proteomic and metabolomic data.

In this application, each subject was treated as a cluster, and the observations obtained at multiple time points from the same subject constituted repeated measurements within that cluster. The proteomic data were represented by ***X*** 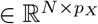, and the metabolomic data were represented by 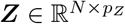, where *p*_*X*_ = 1322 and *p*_*Z*_ = 3529. DOS was used as the response variable ***y*** ∈ ℝ^*N*^ . The dataset contained 150 observations from 53 clusters. Because multiple observations were obtained from the same subject, observations within each subject were expected to be correlated.

For this application, we specified **Φ** = ***AA***^*T*^, where ***A*** ∈ ℝ^*N ×m*^ is the subject-membership matrix whose (*i, k*)th element is one if observation *i* was obtained from subject (cluster) *k*, and zero otherwise. Thus, observations from the same subject shared a common random effect, corresponding to an exchangeable within-subject dependence structure.

Ding et al. [1] analyzed these observations using cooperative learning under the assumption that all observations were independent and identically distributed. In the present analysis, we applied CL-pLMM to account for the correlation among repeated measurements from the same subject.

### 5.2 Evaluation of prediction performance

CL-pLMM was compared with the same methods considered in the simulation study. As in the simulation study, the data were divided into training and test sets at the subject level so that observations from the same subject did not appear in both sets. Ten-fold cross-validation was used for model tuning, and prediction performance was evaluated using the test mean squared error (MSE). Tuning parameters were selected in the same manner as in the simulation study, with the candidate values of the agreement parameter specified as {0, 0.03, 0.05, 0.1, 0.2, 0.4, 0.6, 0.8, 1}.

### 5.3 Results

Tab. 1 presents the test MSE and the number of selected features for each method. Among all the methods considered, CL-pLMM achieved the lowest mean test MSE. This result suggests that jointly using the proteomic and metabolomic data while accounting for the correlation among repeated measurements from the same subject may improve the prediction of DOS.

Compared with the original cooperative learning method, CL-pLMM substantially reduced the mean test MSE from 515.77 to 336.86. Furthermore, CL-pLMM outperformed the methods that used either the proteomic or metabolomic view alone. These findings suggest that the two data views may contain complementary information relevant to the prediction of DOS.

Early Fusion also achieved a relatively low mean test MSE of 349.47, and the difference in mean test MSE between Early Fusion and CL-pLMM was modest. However, the standard deviation of the test MSE was smaller for CL-pLMM than for Early Fusion. This result suggests that CL-pLMM may provide more stable prediction performance across different data splits, in addition to its lower mean prediction error.

**Table 1:** Test MSE and the number of selected features (mean ± SD) on the labor onset dataset.

| Method | Test MSE (mean ± SD) | Selected features (mean ± SD) |
| --- | --- | --- |
| Separate X | 523.22 ± 382.98 | 40.30 ± 18.34 |
| Separate Z | 407.41 ± 89.43 | 37.00 ± 10.59 |
| Early Fusion | 349.47 ± 90.36 | 58.80 ± 28.11 |
| Late Fusion | 455.00 ± 127.99 | 74.90 ± 27.63 |
| Cooperative Learning | 515.77 ± 387.77 | 88.30 ± 49.53 |
| Separate X-pLMM | 477.51 ± 111.63 | 25.10 ± 10.16 |
| Separate Z-pLMM | 401.27 ± 115.92 | 35.60 ± 10.74 |
| Early Fusion-pLMM | 402.19 ± 112.68 | 39.00 ± 16.63 |
| Late Fusion-pLMM | 442.28 ± 178.48 | 59.10 ± 16.60 |
| CL-pLMM | 336.86 ± 78.19 | 70.00 ± 22.15 |

The mean number of features selected by CL-pLMM was smaller than that selected by the original cooperative learning method. Thus, accounting for within-subject correlation may contribute not only to improved prediction performance but also to a more parsimonious feature selection. Nevertheless, CL-pLMM selected more features than Separate X, Separate Z, and Early Fusion. Therefore, CL-pLMM was not uniformly sparser than all the competing methods.

Finally, the dataset contained a total of 4,851 explanatory variables but only 150 observations from 53 clusters. Because the number of variables was considerably larger than the number of observations and clusters, both the selected features and the estimated prediction performance may be sensitive to the particular division of the data into training and test sets.

## 6 Conclusion

In this study, we proposed CL-pLMM, which extends cooperative learning to a penalized linear mixed-effects model for high-dimensional multiview data with a clustered structure. CL-pLMM retains the key feature of cooperative learning by encouraging agreement between the predictions obtained from individual views, while accounting for dependence among observations within the same cluster through a mixed-effects model and performing variable selection using a Lasso penalty. Furthermore, we showed that the proposed objective function can be expressed as a penalized linear mixed-effects model applied to augmented data, thereby allowing existing estimation frameworks to be used.

In the simulation study, when both views contained comparable amounts of predictive information, CL-pLMM achieved the lowest prediction error across all levels of within-cluster dependence considered, and its improvement over standard cooperative learning became greater as the dependence increased. In contrast, when the signal contributions were asymmetric across the views or only one view contained signal, a method based on a single view sometimes performed best. These results suggest that the advantages of CL-pLMM are particularly evident when multiple views contain shared or complementary predictive information. In the real-data analysis, CL-pLMM also achieved the lowest mean test MSE among the methods compared.

Although CL-pLMM is developed for multiview data, the present study considered only two data views. The proposed framework can be extended to more than two views by generalizing the agreement penalty following Ding et al. [1]. However, the performance of such an extension has not yet been investigated, and its evaluation through simulation studies and applications to real-world data remains an important topic for future research.

Regarding the dependence structure, CL-pLMM is formulated for a prespecified positive semidefinite matrix **Φ**, the present study examined only exchangeable within-cluster dependence structures. Specifically, we used 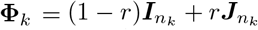 in the simulation study and **Φ** = ***AA***^*T*^, where ***A*** represents the cluster-membership matrix in the real-data analysis. Both specifications of **Φ** assume an exchangeable within-cluster dependence structure, under which all pairs of observations within the same cluster have the same dependence regardless of their measurement times or temporal separation. Therefore, the results of this study do not establish the performance of CL-pLMM under more general dependence structures. Although the formulation can accommodate other prespecified positive semidefinite matrices, further studies are needed to evaluate its prediction and variable-selection performance under structures such as autoregressive, time-distance-based, or unstructured within-cluster dependence.

In the real-data analysis, the dataset was limited to 150 observations from 53 subjects, despite containing 4,851 explanatory variables. In such a small-sample, high-dimensional setting with *p* ≫ *N*, the estimated prediction performance and selected variables may be sensitive to the particular division of the data into training and test sets. Future studies should therefore evaluate the prediction performance and variable-selection stability of the proposed method using datasets containing larger numbers of subjects. Validation using an independent external dataset is also needed to assess the reproducibility of the selected variables and the generalizability of the prediction performance.

The relative prediction performance of the methods in the real-data analysis did not necessarily agree with the results reported by Ding et al. [1]. For example, standard cooperative learning did not outperform some of the competing methods in the present study. This discrepancy may be attributable to differences in the analysis procedure. In particular, we did not perform the variable screening used by Ding et al. [1] during preprocessing. In addition, the training–test splits and cross-validation folds were constructed at the cluster level so that observations from the same cluster were not assigned to different data subsets.

